# Inside-out tracking and projection mapping for robot-assisted transcranial magnetic stimulation

**DOI:** 10.1101/2022.01.30.478399

**Authors:** Yihao Liu, Shuya (Joshua) Liu, Shahriar Sefati, Jing Tian, Amir Kheradmand, Mehran Armand

## Abstract

Transcranial Magnetic Stimulation (TMS) is a neurostimulation technique based on the principle of electromagnetic induction of an electric field in the brain with both research and clinical applications. To produce an optimal neuro-modulatory effect, the TMS coil must be placed on the head and oriented accurately with respect to the region of interest within the brain. A robotic method can enhance the accuracy and facilitate the procedure for TMS coil placement. This work presents two system improvements for robot-assisted TMS (RA-TMS) application. Previous systems have used outside-in tracking method where a stationary external infrared (IR) tracker is used as a reference point to track the head and TMS coil position. This method is prone to losing track of the coil or the head if the IR camera is blocked by the robotic arm during its motion. To address this issue, we implemented an inside-out tracking method by mounting a portable IR camera on the robot end-effector. This method guarantees that the line of sight of the IR camera is not obscured by the robotic arm at any time during its motion. We also integrated a portable projection mapping device (PPMD) into the RA-TMS system to provide visual guidance during TMS application. PPMD can track the head via an IR tracker, and can project a planned contact point of the TMS coil on the head or overlay the underlying brain anatomy in real-time.

## 1. INTRODUCTION

Transcranial magnetic stimulation (TMS) is a safe neuro-modulation technique based on electromagnetic induction of an electric field inside the brain.^1, 2^ TMS has behavioral effects and therapeutic potential and can be used as a valuable tool to probe brain function.^3^ The working principle of TMS is based on its modulatory effect on neural activity induced by the parameters of magnetic stimulation.^1, 3, 4^

To apply TMS and produce an optimal modulatory effect at a target location within the brain, the TMS coil must be placed accurately on the subject’s head.^5, 6^ For this purpose, TMS systems commonly use neuronavigation systems to aid with real-time alignment of the TMS coil on the head. The neuronavigation systems use an IR camera to provide a real-time pose of the TMS coil in six degrees of freedom (6DoF). A pointer with reflective markers is used to obtain a series of landmarks on the head and to register with the corresponding points on the imaging of the head (e.g., MRI). The TMS coil can then be placed and aligned by an operator with the help of an interactive navigation system that provides the 6D poses of TMS coil relative to the head and the brain in real time. Such alignment is done using a crosshair visual guide in conventional TMS system.^7^

In this process, the manual alignment of the coil is inherently inaccurate as it requires a high-level of eye-hand coordination while the operator has to continuously look back and forth between the TMS coil and a display screen of the navigation system. Therefore, the Robot-Assisted TMS (RA-TMS) system is valuable to improve TMS outcomes as it can (i) facilitate the placement of the TMS coil on the head relative to the target location inside the brain, and (ii) accurately maintain and adjust the position and orientation of the TMS coil on the head during magnetic stimulation.^6, 8–13^

The RA-TMS system usually consists of a command computer, a robot manipulator and an optical tracker as shown in Figure 1B. At least two sets of rigid body markers are needed; one to track the head and another one to track the TMS coil or the base of the robot arm. The existing RA-TMS systems, as reported in the literature,^6, 8–13^ use an ‘outside-in tracking’ method in which a stationary, external IR camera is used to track the 6DoF pose of the TMS coil on the robot arm and the subject’s head. In this method, a set of rigid-body markers are attached to the end-effector which carries the TMS coil. This method is, however, prone to loss of tracking of either the head or the TMS coil, as the robot arm may obscure the line of sight of the camera while it is moving towards the target location. To address this limitation, we implement an ‘inside-out’ tracking method where, instead of a fixed external IR camera, a portable IR camera is mounted on the end-effector of the robot, and through mechanical design, the rigid body markers on the head can be always visible to the tracker (Figure 1A and 4). For this purpose, we used a portable projection mapping device (PPMD), previously developed by our group.^14^ PPMD is a compact, portable system, and contains IR stereo cameras with a sub-millimeter tracking accuracy that can be used for the inside-out tracking method. PPMD’s IR camera is significantly cheaper than industrial IR camera, with a reduced work range but a comparable accuracy.

**Figure 1.**
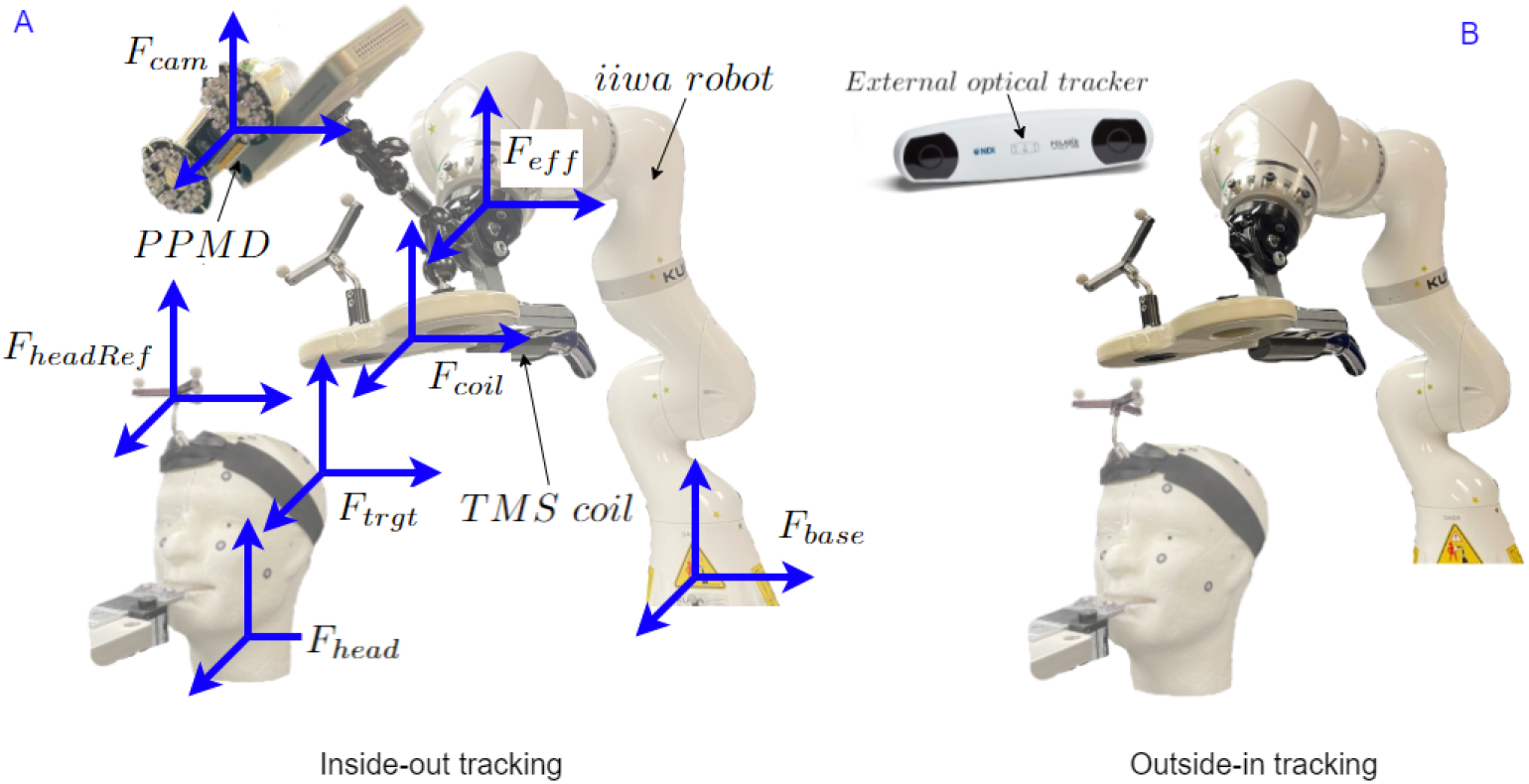
An illustration of inside-out tracking and outside-in tracking for RA-TMS application (note the TMS coil cable is not shown here). Figure A demonstrates the hardware with the key reference frames used in the system. Figure B demonstrates the outside-in tracking method in which the robot arm may block the line of view of IR camera when it moves. Here *cam* denotes the IR camera of PPMD, *headRef* denotes the rigid body marker on the head, *trgt* denotes the target location for coil placement on the head, *coil* denotes the center point of the TMS coil, *head* denotes the MRI coordinate space, and *eff* denotes the end-effector of the robot.

Additionally, as an augmented reality device, PPMD can overlay graphics and videos on surfaces of interest.^14, 15^ For medical applications, PPMD can overlay customized geometries such as anatomical models to tracked objects through a portable laser projector. For TMS application, PPMD can be used to render a planned contact point for the TMS coil and overlay the underlying brain anatomy on the subject’s head. Therefore, the operator can verify the location of the TMS coil on the head along with the target area inside the brain.

Overall, the present study has the following contributions to RA-TMS: 1) an inexpensive inside-out tracking method to improve placement and alignment of the TMS coil on the head, 2) a projection mapping system to visualize the contact point of the TMS coil and underlying brain anatomy, along with 3) the complete workflow for calibration of the proposed RA-TMS system.

## 2. METHOD

The hardware necessary for our proposed system consists of a PPMD system, a Kuka IBR7 iiwa collaborative robot (KUKA AG, Augsburg, Germany), and a computer that runs Ubuntu 20.04. In this work we use an additional external IR camera, Polaris Vicra (Northern Digital, Waterloo, Canada), for system calibration and head registration, however, the proposed system calibration can be eventually performed by using the PPMD. The additional IR camera simplifies the system calibration as it eliminates the need to mount and remount PPMD’s IR camera. The PPMD system includes two parts; an optical tracker made from Intel RealSense D455 camera (Intel Corporation, Santa Clara, USA) and a Laser Beam Pro C200 (KDCUSA, Los Angeles, USA) projector. The calibration methods for PPMD camera and projector is detailed by Liu *et al*.^14^ The system uses the open-source 3D Slicer,^16^ which allows the operator to select target points and registration landmarks on an MRI image. These points and landmarks can then be exported to the robot command controller to compute the desired robot pose. The robot control interface is implemented in ROS and is built based on an iiwa controller implementation by Safeea *et al*.^17^

The setup of the system is illustrated in Figure 1A and 4. In this paper, a reference frame is denoted by *F*_*ref*_ where the subscript *ref* is the reference; a transformation is denoted by 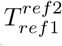 where the superscript *ref*2 is the origin frame and the subscript *ref*1 is the target frame; and a point is denoted by 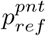 where the subscript *ref* is the reference frame and the superscript *pnt* is the point denotation. For example, *F*_*coil*_ is the frame based on the TMS coil center, and 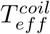 transfers a point 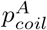 (point A with respect to *F*_*coil*_) to 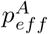 (point A with respect to *F*_*eff*_). The directions of the transformation arrows in all figures are opposite to the directions of the frame conversions, and the arrows show the direction of the transformation matrices. A point set that contains a number of points is denoted by capitalized letter 𝒫.

### 2.1 Workflow

As demonstrated in Figure 2, the workflow for the proposed system consists of steps required to calibrate the coil and PPMD (Section 2.5), register the subject’s head to the head MRI model (Section 2.4), plan a target location for TMS stimulation on the MRI model of the head, overlay the TMS contact point on the head by projection mapping, confirm the target location by visual inspection, and command the robot to align the TMS coil with the target location on the head.

**Figure 2.**
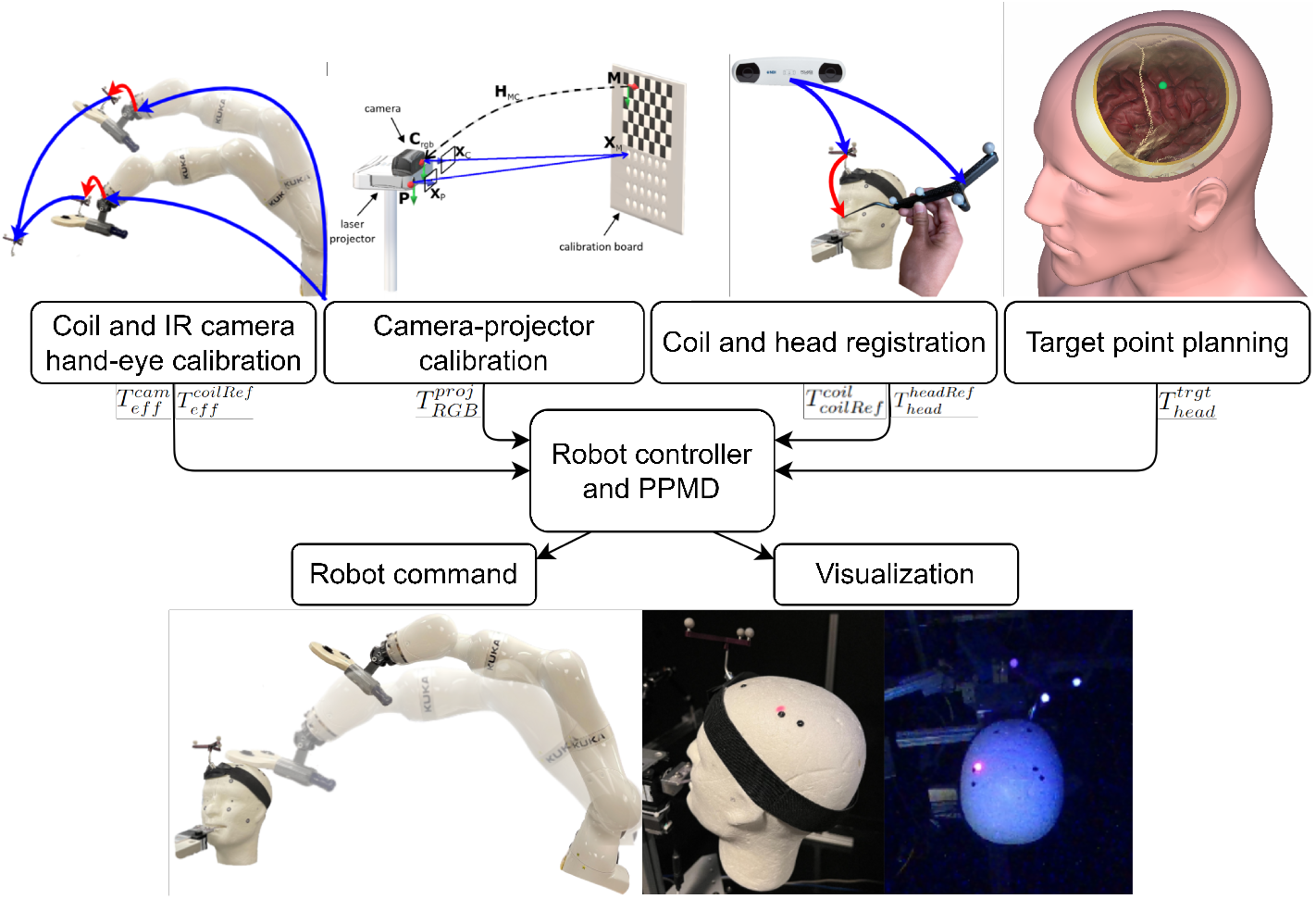
The workflow of the proposed system.

In Figure 3, 4 and 5, calibrations for 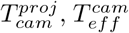 and 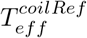, and the registration for 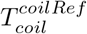 are performed once, as long as the TMS coil, PPMD and the rigid body marker are not moved from where they are attached to. Registration for 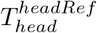 is performed each time if the rigid body marker on the head is moved. A bite bar is used to avoid unwanted head motions during TMS. The accuracy of the system depends on the calibration and registration processes, and since it is hard to determine a ground truth for whether the coil placement is accurate, we designed experiments to individually estimate the accuracy of the transformations that contribute to the total error. The procedures used to evaluate the accuracy of the system are described in Section 3.

**Figure 3.**
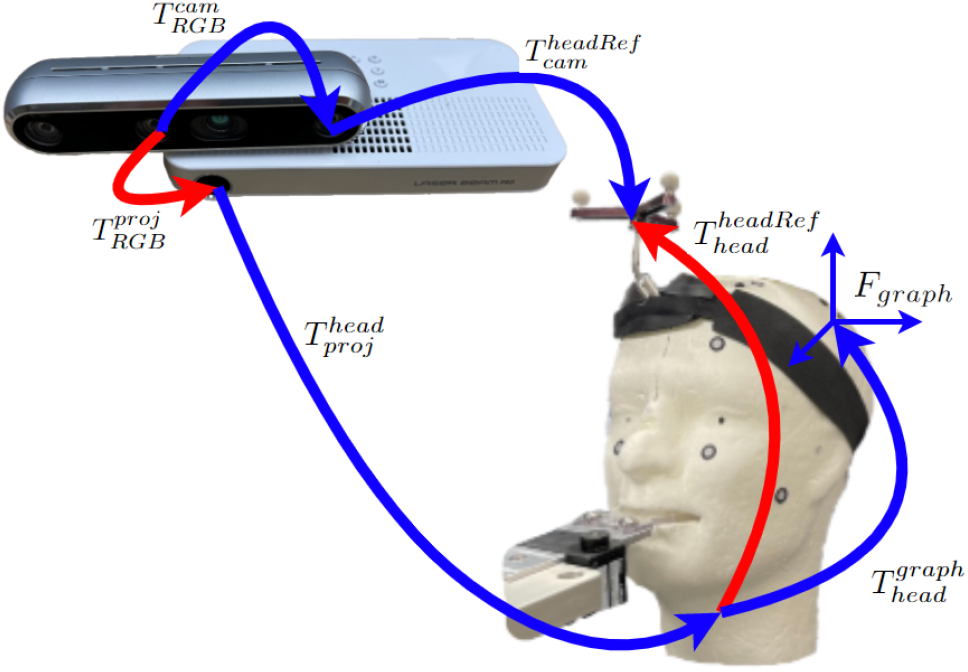
PPMD setup. Red arrows indicate transformations that must be calibrated or registered. 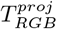 is obtained from the calibration method in,^14^ where *RGB* denotes the reference frame for the RGB camera, and *proj* denotes the reference frame for the projector graphical space. 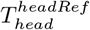 needs to be registered, and the registration method is introduced in Section 2.4. 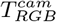 is provided by the manufacturing specification. 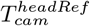 is obtained by optical tracking. *F*_*graph*_ is the reference frame of graphics that needs to be rendered. Projection mapping effect, infrared filters and LEDs are not shown in the image.

**Figure 4.**
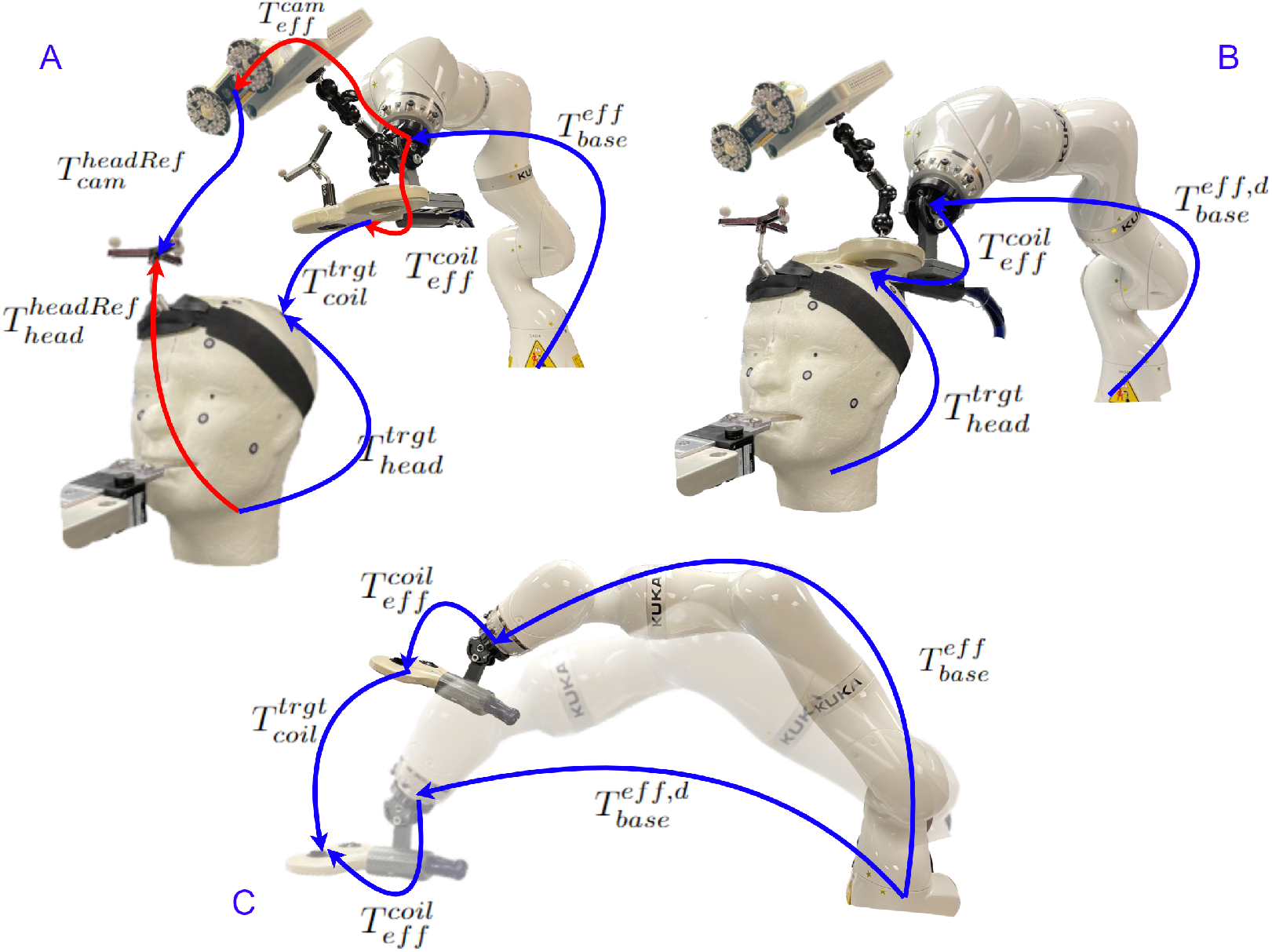
The transformation chain of the key reference frames used for the inside-out tracking method in RA-TMS. Red arrows are the transformations that must be calibrated or registered. Figure A shows the current robot pose and the target TMS location on the head. Figure B shows the desired robot pose when the coil is placed at the target location. Figure C shows 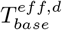 which is derived from the known transformations in A and B. 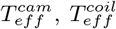 are obtained by calibration, 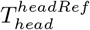 is obtained by registration, 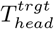 is obtained by planning the target point in 3D Slicer, and 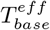 is obtained by the sensors in the robot arm. The arrows show the directions of the transformation matrices and are the opposite to the directions of frame conversions. Note the TMS coil cable is not shown here.

**Figure 5.**
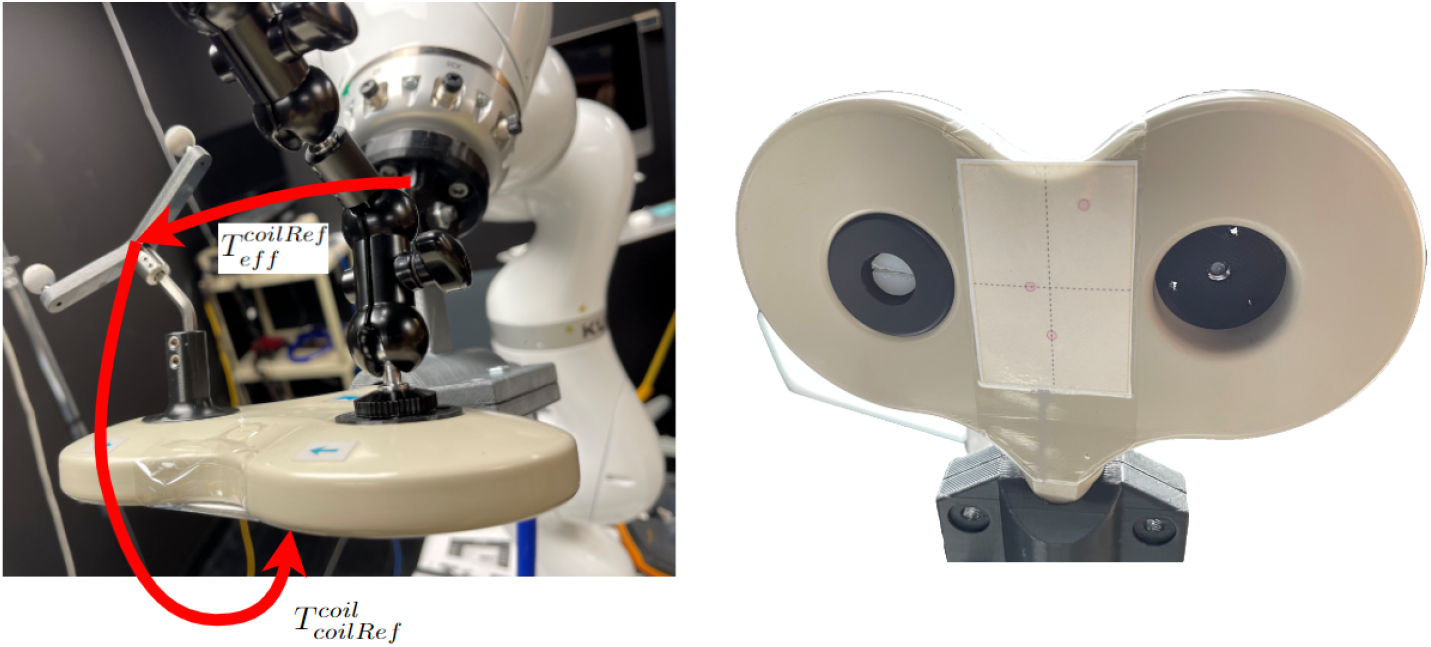
The TMS coil used in the RA-TMS system. Left image shows the transformations used for the calibration of the TMS coil to obtain the transformation from end-effector to the TMS coil 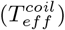. Right image shows the sticker with 12 known coordinates that are used to obtain the transformation from the rigid body marker on the TMS coil to the center of the TMS coil 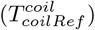

### 2.2 PPMD

PPMD was first developed as an augmented reality device to provide visual assistance during surgery.^14^ For RA-TMS application, PPMD can be used to project planned contact point of the TMS coil and underlying brain anatomy. PPMD uses RealSense D455 which has an RGB camera and two IR cameras to capture binocular infrared reflections of the retro-reflective spheres. The infrared reflections are gray scale images that can be processed to compute the 6D pose of the rigid body marker. As illustrated in Figure 3, PPMD defines the center of the left infrared camera as the *F*_*cam*_ (perspective is from the camera facing out, as defined by the manufacturer), and the center of the projector emitter as the projector space origin *F*_*proj*_. The 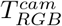 is provided by the manufacturer. Transformation 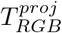 can be obtained by the calibration method detailed in Liu *et al*, ^14^ and 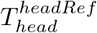 registration is described in Section 2.4.

When projecting planned graphics, the transformation 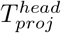 is derived from 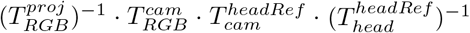. Then, the transformation for the planned graphics with respect to the projector space 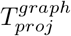 can be obtained by 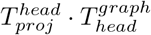. These transformations are displayed in Figure 3.

### 2.3 Inside-out Tracking

To apply inside-out tracking of the robot arm, a number of transformations must be computed. These transformations are illustrated in Figure 4 where 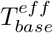 is the current robot pose (Figure 4A) and 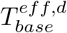 is the desired robot pose (Figure 4B). Accordingly, 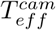 is the transformation from the end-effector of the robot to the left IR imager of PPMD; 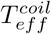 is the transformation from the end-effector to the center of the TMS coil; 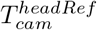 is the transformation from the left IR imager of PPMD to the rigid body marker on the head; 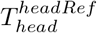 is the transformation from the head center to the rigid body marker on the head (obtained from the registration of the MRI model to the subject’s head as described in Section 2.4); 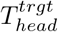 is the transformation from the head coordinate system (also the MRI coordinate system) to the TMS target location on the head; and 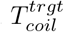 is the transformation from the TMS coil to the TMS target location.

The goal is to derive 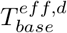 that minimizes 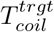 to an identity matrix (Figure 4). The derivation of 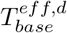 is shown in Equation 1, 2 and 3. On the right hand side of the Equation 1, 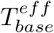 can be obtained from robot controller; 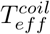 can be obtained by the calibration of the coil as described in Section 2.5; and 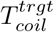 can be derived from Equation 2, using transformations illustrated in Figure 4A. On the right hand side of the Equation 2, 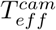 is obtained by the hand-eye calibration of the IR camera (Section 2.5); 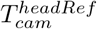 is obtained by the IR camera of PPMD; 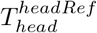 is obtained by registering the MRI image to subject’s head, as described in Section 2.4; and 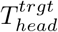 is obtained by identifying the target location on the MRI to apply TMS. Combining Equation 1 and 2, the calculation can derive 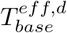, which is the desired pose of robot arm that is sent to the robot controller.

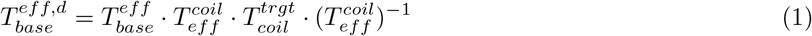

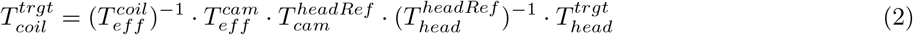

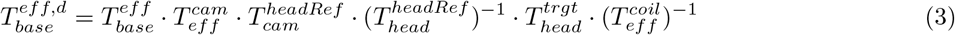

### 2.4 Registration

The target location for TMS coil placement on the head, 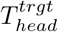, can be selected on the MRI model. To obtain 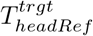, which is the target location with respect to the coordinate system of the rigid body marker on the head, we need to obtain the transformation from the MRI model to the rigid body marker on the head, 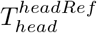, using pair-point registration.^18^ For this purpose, a pair of point sets are selected; one set on the head 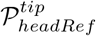 and the corresponding set on the MRI, 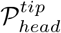. These corresponding points are unambiguous anatomical landmarks such as the tip of the nose, the tragus or the nasion which can each be identified and accurately marked on both the MRI image and the subject’s head. In this paper, these points are also referred to as ‘fiducials’.

To perform the pair-point registration, the operator uses a tracked pointer to collect the coordinates of the anatomical landmarks on the head. The collection process, termed ‘digitization’, is done by touching the landmarks using the tracked pointer. The pointer tip coordinate is then transformed to the coordinate frame of the rigid body marker on the head, which is also tracked by the IR camera to obtain 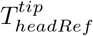. Taking only the translation part in 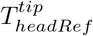, the point set 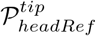 is then obtained. The other set 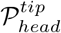 is selected using the corresponding landmarks from the MRI.

### 2.5 Calibration

In order to derive the transformation from the end-effector to an attached location, a hand-eye calibration or accurate modeling of the mechanical part is performed.^19, 20^ Because accurate models of the TMS coil and the mechanical part attached to the end-effector were not available, we performed hand-eye calibration to derive 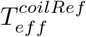 and 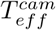. The hand-eye calibration is modeled by an *AX* = *XB* problem, where *X* is the unknown transformation. The transformation from the robot end-effector to the center of the TMS coil can be broken down into two parts, as shown in Figure 5A: (i) 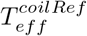 as the transformation from the robot end-effector to the rigid body marker mounted on the TMS coil, and (ii) 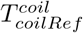 as the transformation from the rigid body marker to the TMS coil center. 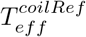 is the *X* frame in the *AX* = *XB* and is solved in the following equations:

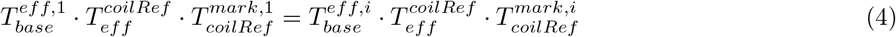

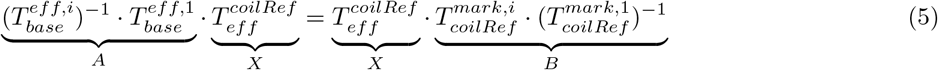

*F*_*mark*_ is an additional rigid body marker at a fixed distance from the robot that is used in the calibration process. During the calibration process (Figure 6), the operator moves the robot to different poses to collects data for the corresponding 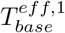 to 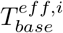 and 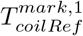 to 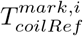. The *AX* = *XB* is then solved by the optimization method proposed by Park *et al*.^21^

**Figure 6.**
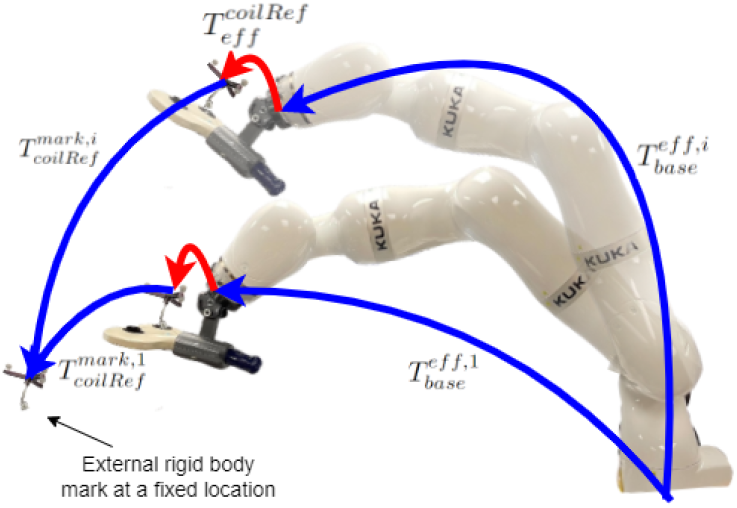
Hand-eye calibration for the transformation from the end-effector to the rigid body marker attached on the TMS coil 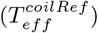. The reference frame *F*_*mark*_ is a stationary rigid body marker. Red arrows are the transformation to be obtained by the hand-eye calibration. 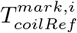 are obtained by IR camera, and 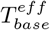 are obtained by robot sensors.

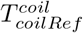 can be solved by the pair-point registration process described in Section 2.4. A sticker with 12 known coordinate points is attached to the surface under the TMS coil, with its center aligned with the coil center. Point set 𝒫_*coilRef*_, which is a set of points with respect to *F*_*coilRef*_, are collected using the tracked pointer. Then, 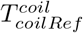 is registered using the same pair-point registration method described in Section 2.4. Finally, the transformation 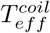 is obtained by 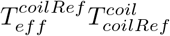.

Similar to 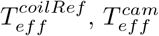 is obtained using hand-eye calibration by setting up the *AX* = *XB* problem.

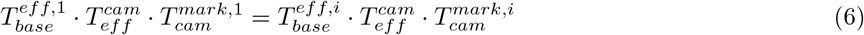

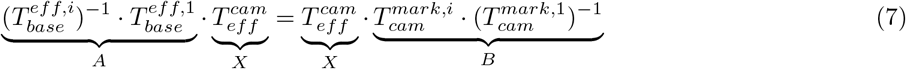

## 3. EXPERIMENTS

We performed experiments to evaluate accuracy of the system. In the process of the registration and calibration, 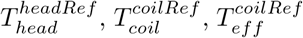 and 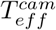 are the transformations that contributed to the errors of calculated robot pose and the projection mapping by PPMD. Here we examined the accuracy of these transformations. Note these errors are in addition to the PPMD tracking error (error of 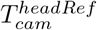), which was previously validated by Liu *et al*.^14^ Additionally, the accuracy of robot pose 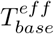 is dependent on the specification of the iiwa robot, and 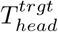 is considered to be accurate since it depends on just locating a target TMS location on the MRI. We used a laser-scanned foam head model in place of MRI. The foam head was attached with a number of 3D printed targets, which can be easily identified in the laser-scanned model.

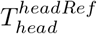 and 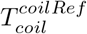 are obtained by the pair-point registration. For 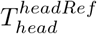, a set of known points in MRI coordinate system *F*_*head*_, are paired with the corresponding points with respect to the frame *F*_*headRef*_. The point set 𝒫_*headRef*_ is collected by digitizing the landmarks using a tracked pointer. Similarly, for 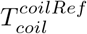, a set of known points on the sticker attached to the TMS coil are paired with the corresponding points with respect to the frame *F*_*coilRef*_. We used fiducial registration error (FRE) and target registration error (TRE) to validate the registration accuracy per Maurer *et al*.^22^ Registration residuals (Equation 8) measures the difference between the fiducial coordinates and their calculated positions after applying the registration transformation. FRE (Equation 9) is the root mean square of the registration residuals for all fiducial points. TRE (Equation 10) measures the distance between the target location on the head or the TMS coil center and their calculated positions after applying 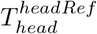 and 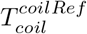, respectively.

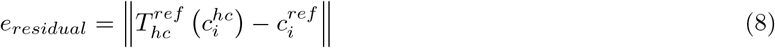

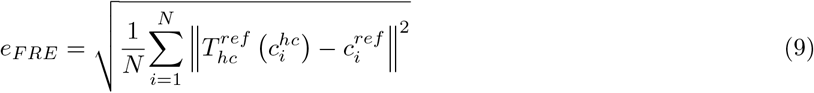

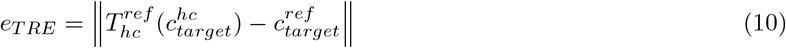

To validate the hand-eye calibration of 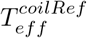 and 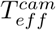, we verified their repeatability (deviation from the mean and standard deviation). In addition, *AX*− *XB* and (*AX*)^−1^*XB* were also verified, and we refer to them as ‘residuals’ in the following sections. Repeatability shows the consistency of hand-eye calibration results across different trials. We performed 16 hand-eye calibrations for both 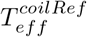 and 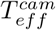. Residuals measure how well the optimization is for each data entry in deriving *X*, and they should be minimized in the optimization algorithm when solving *AX* = *XB*.^21^ For each data entry collected in Equation 5 and 7, a residual error is calculated by deriving the discrepancy between the left hand side and right hand side of the equation. The smaller the resultant translation and rotation, the more effective the optimization is in solving the *AX* = *XB* problem.

**Figure 7.**
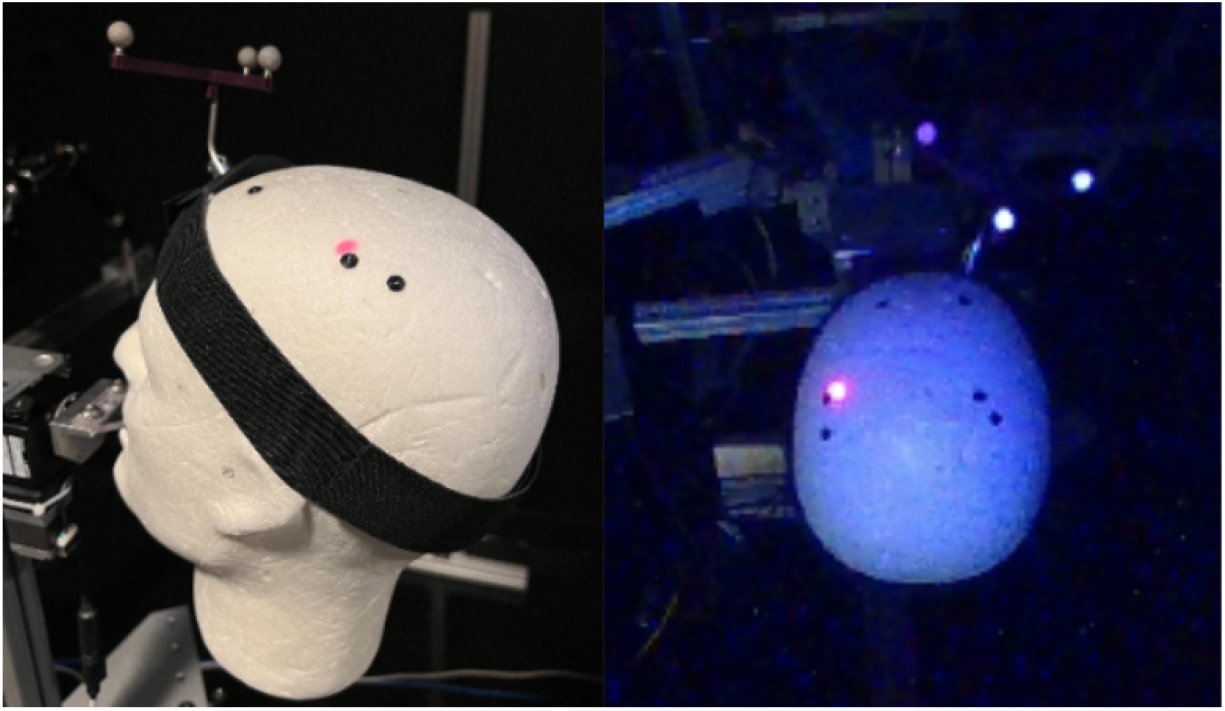
Projection of a red dot on the head by PPMD to mark the planned target location for TMS coil placement. These target locations are also marked by a black circular sticker on the head. Left image is the user view, and right image is the view from PPMD RGB camera.

## 4. RESULTS

### 4.1 FRE and Residual Error of TMS Coil Registration

For the registration of the TMS coil, we used a tracked pointer to touch the 12 points on the sticker attached to the TMS coil (Figure 5). The digitized coordinates are paired with the known coordinates on the sticker, by which 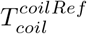 is derived. The process was repeated six times and the results are shown in Table 1. Each row for fiducial point number 1 to 12 shows the residual errors after applying the registration transformation. The center of the coil (row 12) is also used as a fiducial, so its residual error is not TRE by the formal definition, but can be used as an estimation of TRE. FRE is shown in the last row. In the table, each of the columns 1 to 6 corresponds to one registration result. These results demonstrate that the registration errors for the TMS coil were trivial, with residual values and FRE values less than 1 mm, which are consistent across different trials.

**Table 1.** Residual error at each fiducial and FRE for the registration of the TMS coil 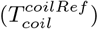. All units are in millimeters. Fiducial 12 is the center of the coil, and residual at fiducial 12 is an estimate of TRE.

| Fiducial number | Trial number |  |  |  |  |  |  |  |
| --- | --- | --- | --- | --- | --- | --- | --- | --- |
|  | 1 | 2 | 3 | 4 | 5 | 6 | SD | Mean |
| 1 | 0.5011 | 0.6502 | 0.3598 | 0.9246 | 0.9066 | 0.8941 | 0.2401 | 0.7061 |
| 2 | 0.3451 | 0.3721 | 0.5499 | 0.1439 | 0.3125 | 0.5705 | 0.1592 | 0.3823 |
| 3 | 0.3525 | 0.1552 | 0.3314 | 0.4638 | 0.4010 | 0.3024 | 0.1045 | 0.3344 |
| 4 | 0.6082 | 0.5167 | 0.4792 | 0.3461 | 0.2303 | 0.3682 | 0.1358 | 0.4248 |
| 5 | 0.3203 | 1.1796 | 0.7871 | 0.3319 | 0.7401 | 0.2999 | 0.3550 | 0.6098 |
| 6 | 0.2395 | 0.3661 | 1.0249 | 0.7995 | 0.6397 | 0.5475 | 0.2861 | 0.6029 |
| 7 | 0.5807 | 0.4336 | 0.7623 | 0.5678 | 0.4389 | 0.3678 | 0.1426 | 0.5252 |
| 8 | 0.7773 | 1.0985 | 0.6166 | 0.3699 | 0.5512 | 0.3605 | 0.2785 | 0.6290 |
| 9 | 0.4255 | 0.4353 | 0.5344 | 0.5701 | 0.6786 | 0.2356 | 0.1518 | 0.4799 |
| 10 | 0.1588 | 0.3274 | 0.3417 | 0.3614 | 0.4667 | 0.3588 | 0.0998 | 0.3358 |
| 11 | 0.1594 | 0.3612 | 0.2848 | 0.8031 | 0.4913 | 0.5577 | 0.2269 | 0.4429 |
| 12 (center) | 0.1601 | 0.2045 | 0.0511 | 0.3749 | 0.1094 | 0.2833 | 0.1179 | 0.1972 |
| FRE | 0.4304 | 0.5945 | 0.5693 | 0.5523 | 0.5415 | 0.4639 | 0.0640 | 0.5253 |

### 4.2 TRE and FRE for Head Registration

To estimate the registration accuracy, we performed the registration for the foam head five times per the definition of TRE in Section 3. Six fiducials were used for the registration, and four different TMS targets were labelled on the foam head and its laser-scanned model. The coordinates of these four target locations were used as the ground truth to derive TRE. Table 2 presents the TRE of the registrations where each column is one registration trial and each row is the results for one of the four targets. Table 3 presents the residual and FRE for each of the five head registration trials. TRE of the four targets was within the range of 1 to 2.5 mm, and the average FRE on the six selected fiducials was 2.5229 mm. TRE and FRE of head registration are expected to be larger than the coil registration, because the landmarks used as fiducials are not marked on the head so they are harder to identify as opposed to the ones used for coil registration (Section 5).

**Table 2.**
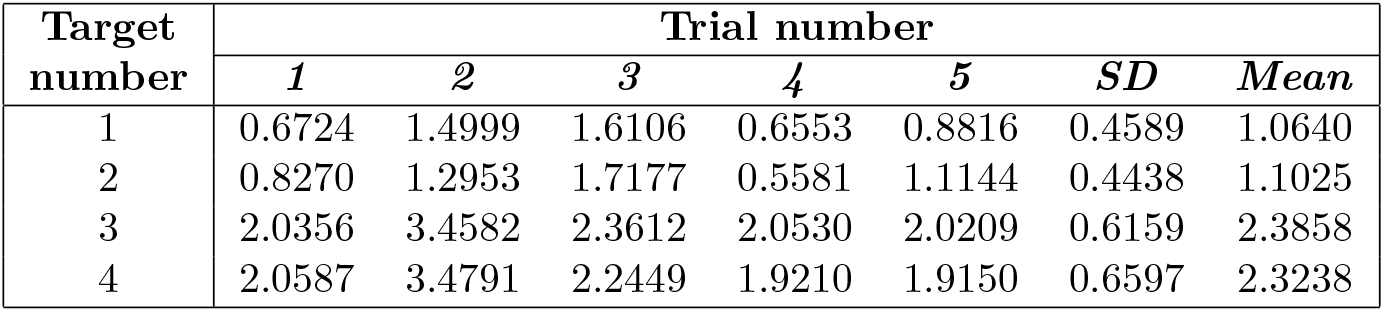
TRE at each target point for the head registration 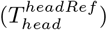. All units are in millimeters.

| Target number | Trial number |  |  |  |  |  |  |
| --- | --- | --- | --- | --- | --- | --- | --- |
|  | 1 | 2 | 3 | 4 | 5 | SD | Mean |
| 1 | 0.6724 | 1.4999 | 1.6106 | 0.6553 | 0.8816 | 0.4589 | 1.0640 |
| 2 | 0.8270 | 1.2953 | 1.7177 | 0.5581 | 1.1144 | 0.4438 | 1.1025 |
| 3 | 2.0356 | 3.4582 | 2.3612 | 2.0530 | 2.0209 | 0.6159 | 2.3858 |
| 4 | 2.0587 | 3.4791 | 2.2449 | 1.9210 | 1.9150 | 0.6597 | 2.3238 |

**Table 3.** Residual error at fiducials (landmarks) and FRE for the head registration 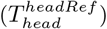. All units are in millimeters.

| Fiducial number | Trial number |  |  |  |  |  |  |
| --- | --- | --- | --- | --- | --- | --- | --- |
|  | 1 | 2 | 3 | 4 | 5 | SD | Mean |
| 1 | 3.6694 | 3.8790 | 3.9550 | 3.5043 | 3.4700 | 0.2174 | 3.6955 |
| 2 | 0.7929 | 1.1996 | 0.6838 | 0.9307 | 1.7203 | 0.4138 | 1.0655 |
| 3 | 2.2546 | 2.1767 | 2.1661 | 1.7918 | 1.4639 | 0.3354 | 1.9706 |
| 4 | 3.6840 | 3.7645 | 3.7011 | 4.3827 | 2.5208 | 0.6747 | 3.6106 |
| 5 | 2.2801 | 1.0522 | 2.2785 | 2.7676 | 2.1257 | 0.6341 | 2.1008 |
| 6 | 1.2789 | 1.6398 | 0.8221 | 0.9948 | 0.7389 | 0.3681 | 1.0949 |
| FRE | 2.5685 | 2.5557 | 2.5938 | 2.7146 | 2.1818 | 0.2008 | 2.5229 |

### 4.3 Repeatability and Residuals of Hand-eye Calibration

The hand-eye calibration was repeated 16 times for both the TMS coil and the PPMD’s IR camera. Each hand-eye calibration collects 270 entries of *AX* = *XB* data. The repeatability is shown in Table 4, where each row is the standard deviation of 16 transformations, divided into translation and rotation components. Repeatability is also shown in Figure 8, where each line is a trial of the hand-eye calibration. The translations and rotations of each line is calculated by subtracting their mean from the 16 translations and rotations for both 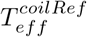 and 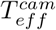. The residuals of *AX* − *XB* and (*AX*)^−1^*XB* are shown in the boxplots in Figure 9 and 10. The calculated residuals (refer to Equation 5 and 7) of each trial are grouped in one box, using both the norms of the translation and rotation vectors. The hand-eye calibration results of the transformation from the end-effector to the TMS coil from Noccaro *et al*.^6^ are also shown for comparison in Figure 9. The standard deviations of translation are around 2 mm and 7 mm for 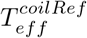 and 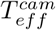 respectively, and the corresponding standard deviations of rotation are around 0.3 degrees and 0.5 degrees. The median of *AX* − *XB* errors were within range of 1-2 mm and 4-8 mm for 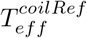 and 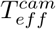, respectively, and the corresponding rotation errors were around 0.1-0.3 degrees and 0.25-0.75 degrees.

**Table 4.** Repeatability of the hand-eye calibration for the transformation from end-effector to the rigid body marker attached to the TMS coil 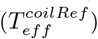 and the transformation from the end-effector to the IR camera of PPMD 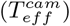, quantified as the standard deviations of 16 trials. *x, y, z* values are for translation and *rx, ry, rz* values are for rotation.

| Transformation | Standard deviation |  |  |  |  |  |
| --- | --- | --- | --- | --- | --- | --- |
| | $x(mm)$ | $y(mm)$ | $z(mm)$ | $rx(deg)$ | $ry(deg)$ | $rz(deg)$ |
| $T_{eff}^{coilRef}$ | 2.4274 | 2.6341 | 1.9052 | 0.3067 | 0.2122 | 0.3387 |
| $T_{eff}^{cam}$ | 7.7603 | 7.9237 | 5.7805 | 0.3507 | 0.8511 | 0.4546 |

**Figure 8.**
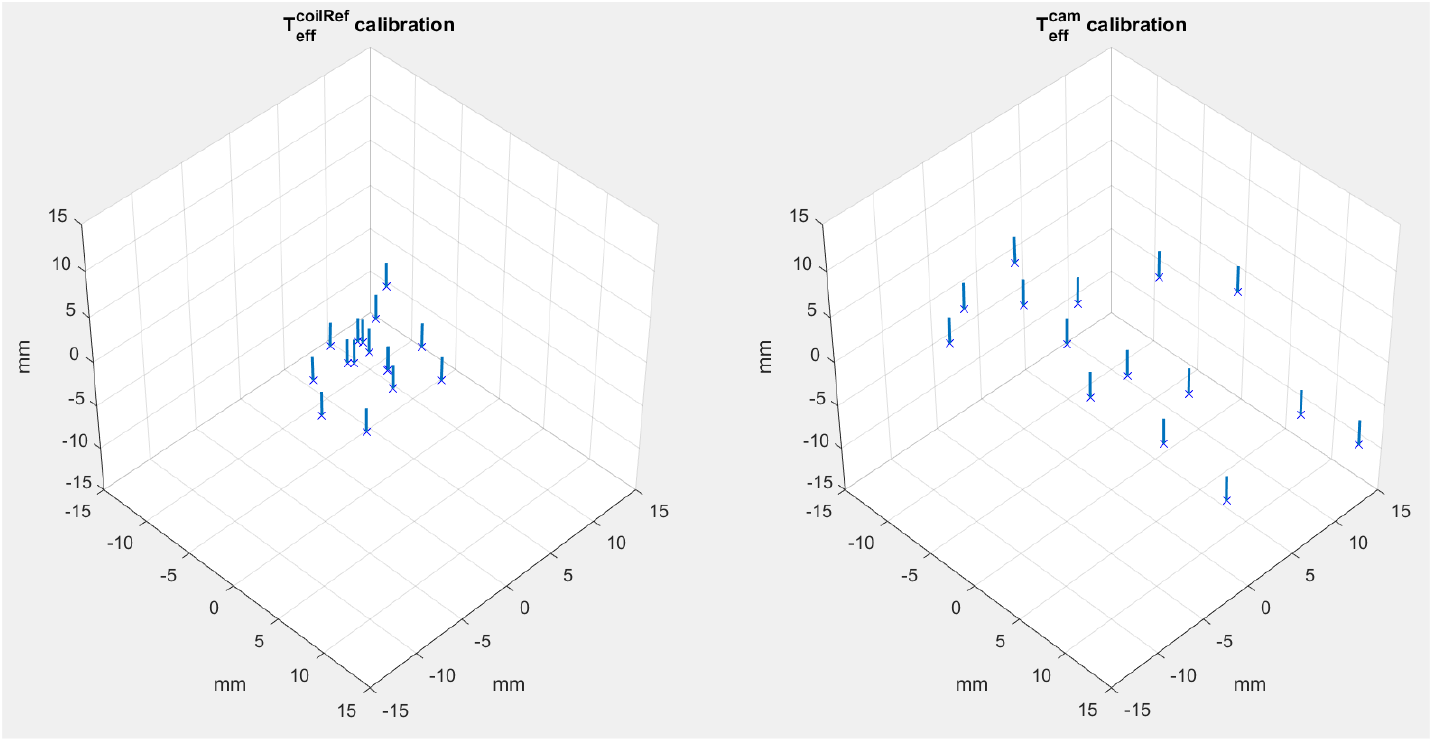
Repeatability of the hand-eye calibration for the transformation from end-effector to the rigid body marker attached to the TMS coil 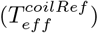 [left] and the transformation from the end-effector to the IR camera of PPMD 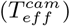 [right]. Each trial of hand-eye calibration is shown as a line with the position and orientation of the 16 lines are calculated by the deviation from the mean of all trials. One line was an outlier and was not shown in the images. Axes units are in millimeter.

**Figure 9.**
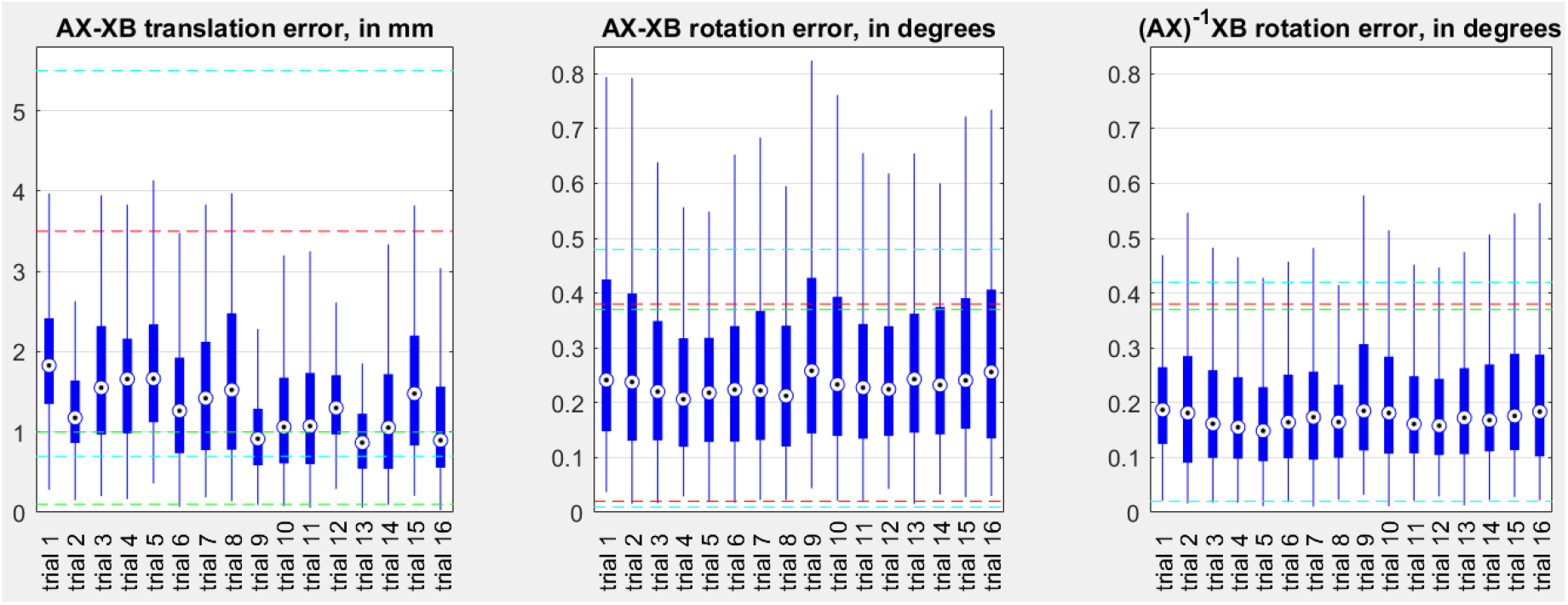
*AX* − *XB* residual and (*AX*)^−1^*XB* residual of hand-eye calibration for the transformation from end-effector to the rigid body marker attached to the TMS coil 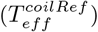. Translation errors of *AX* − *XB* and (*AX*)^−1^*XB* are the same. Each column represents quartiles of 270 entries. Dashed lines are the 75 and 25 percentiles of three different hand-eye calibration methods reported by Noccaro *et al* (estimated from graph in the paper).^6^

**Figure 10.**
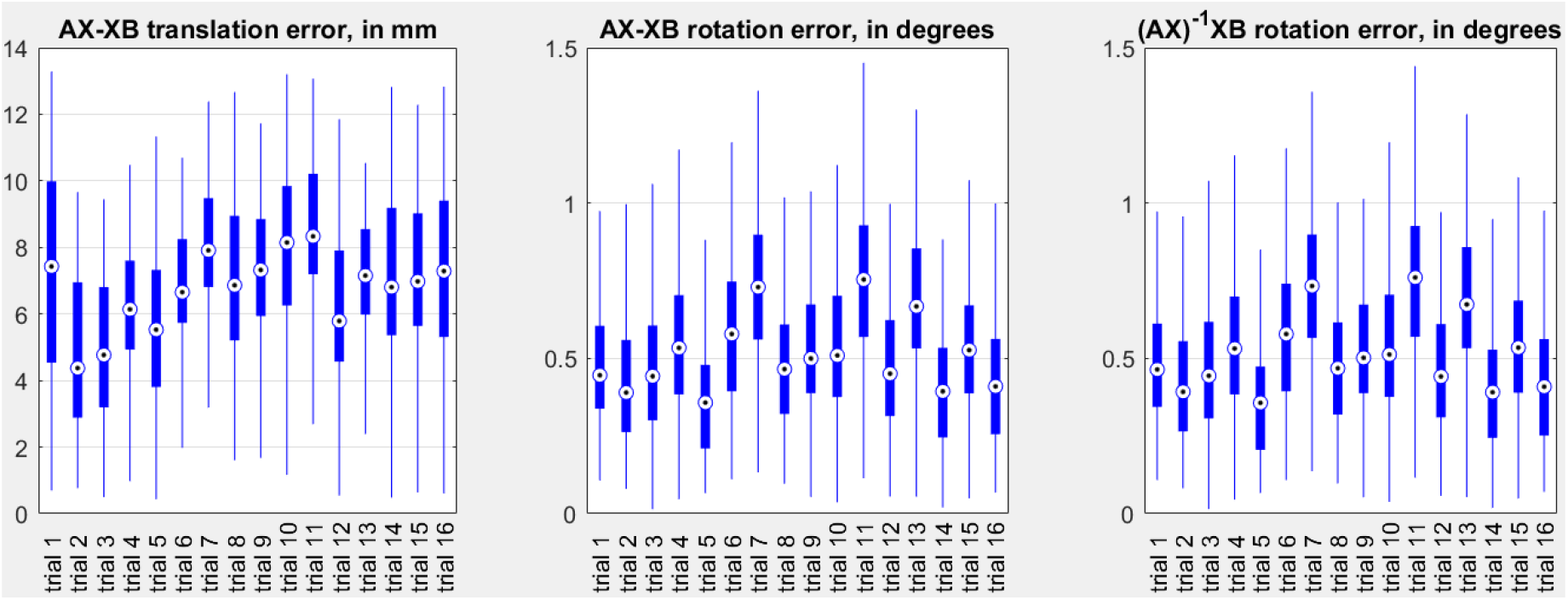
*AX* − *XB* error and (*AX*)^−1^*XB* error of hand-eye calibration for the transformation from the end-effector to the IR camera of PPMD 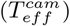. Each column represents the quartiles of 270 data entries.

## 5. DISCUSSION

We have developed an RA-TMS system that improves TMS coil placement on the head by using an inside-out tracking method. Since in this method, the IR camera is mounted on the robot end-effector, and it does not rely on an external IR camera for tracking, the robot arm cannot block the camera and disrupt tracking during its motion. Here we also incorporated PPMD into the RA-TMS system to facilitate visual guidance for TMS placement and alignment on the head. PPMD can track the 6D pose of the head and project graphics to mark a planned contact point of the TMS coil or overlay underlying brain anatomy on the head in real time. In addition, using PPMD can also significantly decrease the cost of the system, by having a smaller work range but a comparably accurate performance. We performed experiments to estimate the errors of calibration and registration in the RA-TMS system that contribute to an overall error in TMS coil placement.

The registration of the head is expected to be less accurate than the registration of the TMS coil, as the errors in locating the landmarks on the head are larger than the ones used to register the coil. Examples of such localization error can be seen in Table 3, where the fiducials 1 and 4 have relatively larger residuals than the other fiducials. This is caused by a lower accuracy in localizing the corresponding landmark from the head to the scanned model. The mean of the FRE is 2.5229 mm, which can be further reduced by better selection of the landmarks, i.e. selecting the landmarks that are easier to access, and clearer on the image. In practice, if the registration is performed using MRI, more accurate results can be obtained by affixing fiducials to the subject’s head before taking the MRI, and by keeping them in place until the registration is concluded. An affixed fiducial is easier to identify than an anatomical landmark. However, this is only viable if the MRI and RA-TMS are done within a close time frame.

Because accurate models were not available for the mechanical parts of the robot end effector and the TMS coil, we instead looked at the repeatability of the hand-eye calibration as a measure of the error in 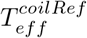. In addition, we verified *AX* − *XB* and (*AX*)^−1^*XB* of the hand-eye calibration data. Our results for the hand-eye calibration of the TMS coil are comparable to Noccaro *et al*, in which the authors defined (*A*_*i*_*X*)^−1^*XB*_*i*_ as the hand-eye calibration error,^6^ and *A*_*i*_*X* − *XB*_*i*_ as the residual. As also shown in previous studies, the hand-eye calibration is prone to low repeatability in TMS application. This is mainly related to inaccurate tracking, inaccurate robot sensors, and unstable robot base or non-rigid mechanical parts that hold the TMS coil. Depending on the optimization algorithm, Richter *et al*. reported variations from around few millimeters to tens of millimeters.^23^ In Richter *et al*., the best repeatability for the hand-eye calibration was derived using a realtime algorithm which modifies the transformation matrix to be non-orthogonal. The non-orthogonal methods relax the constraint of a rigid body rotation matrix and allows more degrees of freedom. However, in Noccaro *et al*, the non-orthogonal method could not improve (*A*_*i*_*X*)^−1^*XB*_*i*_ or *A*_*i*_*X* − *XB*_*i*_ residuals. Figure 8 shows the solution space of the calibrated transformations of our method. As shown by Richter *et al*., the non-orthogonal results tend to gather into a corner of the solution space, while the solutions of conventional hand-eye calibration are more spread. The residuals of non-orthogonal methods are also smaller, because the degrees of freedom of the transformation matrix is larger. Thus, further work is required to improve the hand-eye calibration of the TMS coil for RA-TMS application.

The hand-eye calibration of PPMD’s IR camera depends on its tracking accuracy. The tracking accuracy was reported to be sub-millimeter when the PPMD’s IR camera is less than 20 cm apart from a tracked rigid body marker.^14^ The error increases when the IR camera moves further from the rigid body marker, which is unavoidable in the process of hand-eye calibration, as the robot may move to multiple poses far from the rigid body marker. The PPMD in this study used Intel RealSense D455 for IR tracking, but devices with higher IR imager resolution can be used to improve the accuracy and range of tracking.

## 6. CONCLUSION AND FUTURE WORK

This work introduces an inside-out tracking method for RA-TMS along with a projection mapping technique to visualize the planned TMS target location on the head. We addressed the problem of blocked sight of view of IR camera that can be encountered in outside-in tracking method and improved user experience with a projection mapping technique. Experiments were designed to validate the transformation errors that affect TMS coil placement. Overall, the accuracy of coil placement depends on the hand-eye calibration and registration errors. The coil registration errors are minimal as long as the IR camera has close to sub-millimeter accuracy, but the pair point registration of the head and MRI model can have larger errors related to the accuracy of locating fiducials. Overall, the hand-eye calibration process for the TMS coil is prone to low repeatability and accuracy as also shown in previous RA-TMS systems.^6, 23^ To this end, the accurate coil modeling can be a way to minimize the need for calibration in the future. On the other hand, the hand-eye calibration for the IR camera depends on its tracking accuracy, which can improve by using more accurate IR cameras.

## 7. ACKNOWLEDGMENTS

This work was supported by a grant from the National Institute of Deafness and Other Communication Disorders (R01DC018815). We thank Alejandro Martin-Gomez for comments and assistance with preparing a figure in the manuscript.

